# Engineered orthogonal quorum sensing systems for synthetic gene regulation

**DOI:** 10.1101/499681

**Authors:** Stefan J. Tekel, Christina L. Smith, Brianna Lopez, Amber Mani, Christopher Connot, Xylaan Livingstone, Karmella A. Haynes

**Affiliations:** School of Biological and Health Systems Engineering, Arizona State University, Tempe, AZ; School of Life Sciences, Arizona State University, Tempe, AZ

**Keywords:** Quorum Sensing, Homoserine Lactone, Transcription Factor, Gene Circuit, Biosensor

## Abstract

Gene regulators that are controlled by membrane-permeable compounds called Homoserine lactones (HSLs) have become popular tools for building synthetic gene networks that coordinate behaviors across populations of engineered bacteria. Synthetic HSL-signaling systems are derived from natural DNA and protein elements from microbial quorum signaling pathways. Crosstalk, where a single HSL can activate multiple regulators, can lead to faults in networks composed of parallel signaling pathways. Here, we report an investigation of quorum sensing components to identify synthetic pathways that exhibit little to no crosstalk in liquid and solid cultures. In previous work, we characterized the response of a single regulator (LuxR) to ten distinct HSL-synthase enzymes. Our current study determined the responses of five different regulators (LuxR, LasR, TraR, BjaR, and AubR) to the same set of synthases. We identified two sets of orthogonal synthase-regulator pairs (BjaI/BjaR + EsaI/TraR and LasI/LasR + EsaI/TraR) that show little to no crosstalk when they are expressed in Escherichia coli BL21. These results expand the toolbox of characterized components for engineering microbial communities.

## Introduction

Natural quorum sensing systems control density dependent cell-cell signaling, allowing bacteria to communicate gene expression states through small membrane-permeable molecules. Engineered quorum sensing systems have been used for multiple applications such as biofilm formation (Mukherjee et al., 2017), oscillators (Taylor et al., 2009), pattern formation (Basu et al., 2005), pathogen detection (Jayaraman et al., 2017), synthetic circuits (Wu et al., 2014), and metabolic engineering (Gupta et al., 2017). One class of signalling molecules known as homoserine lactones (HSLs) has been extensively characterized and used for synthetic cell-cell communication. This system is particularly attractive for bioengineering since HSL-signaling components can be used a modules within gene circuits and are portable to non-native strains. Additionally, natural HSL pathways exhibit a biochemical diversity that can be leveraged for circuits controlled by combinations of unique signals (Davis et al., 2015).

A central component of HSL signalling is the synthase enzyme, a protein that catalyzes the formation of HSLs from an acyl carrier protein or coenzyme A and S-adenosyl methionine. We previously characterized a library of synthase enzymes from various bacterial strains in an Escherichia coli host for their ability to generate HSLs that activated a Lux-derived reporter (Daer et al., 2018). This reporter vector was comprised of a constitutively expressed Lux regulator and a GFP expression cassette driven by a lux-box promoter. Lux regulator protein binding to an HSL induces homodimerization and results in binding of the dimer to the lux-box promoter and induction of GFP expression. Our work showed that most of the synthases, which were expected to produce a broad range of short to long-chain HSLs, induced Lux regulator-driven expression to varying degrees. This systematic comparison in a single chassis (E. coli) provided a basis for using diverse synthases for microbial engineering. The synthases that showed little to no stimulation of LuxR might be strong inducers of other regulator orthologues, allowing the operation of distinct HSL signaling pathways in parallel with no crosstalk.

Similar to LuxR, many quorum sensing systems are characterized by a regulator protein and a corresponding promoter that contains a regulator binding motif (e.g. the lux-box from the lux operon in V. fischerii) followed by a weak core promoter motif. Positive quorum sensing regulators (e.g. LuxR) gain affinity for the binding motif after they become bound by HSL ligands. The regulator-HSL complex then recruits RNA polymerase (RNAP) to the core promoter to activate gene expression, as suggested by in vitro studies of LuxR and TraR (Fuqua and Greenberg, 2002). The diversity of quorum sensing regulators can be leveraged to build orthogonal pathways, as was demonstrated by Scott et al who successfully utilized the Tra and Rpa systems with no cross-signaling between inducers and regulators expressed in the same cell. In our report, we describe and characterize the HSL-dependent response of promoter-regulator sets using synthetic HSL ligands and HSLs generated from our E. coli expressed synthases. Scott et al. demonstrated that lux-box-like sequences from four quorum sensing systems (Tra, Las, Lux, Rpa, and Tra) cloned upstream of a universal minimal promoter can be used as HSL-inducible gene regulators (Figure 1) (Scott and Hasty, 2016). Here, we refer to these and similar constructs as “hybrid promoters.” We constructed seven “Receiver” vectors that contain a constitutively expressed HSL-inducible regulators and a corresponding hybrid promoter followed by a GFP reporter. Four included previously tested components from the Las, Lux, Rhl, Rpa, Tra, pathways. Two regulators, derived from the Aub and Bja systems, were tested for the first time as modular circuits in our work.

**Figure 1.**
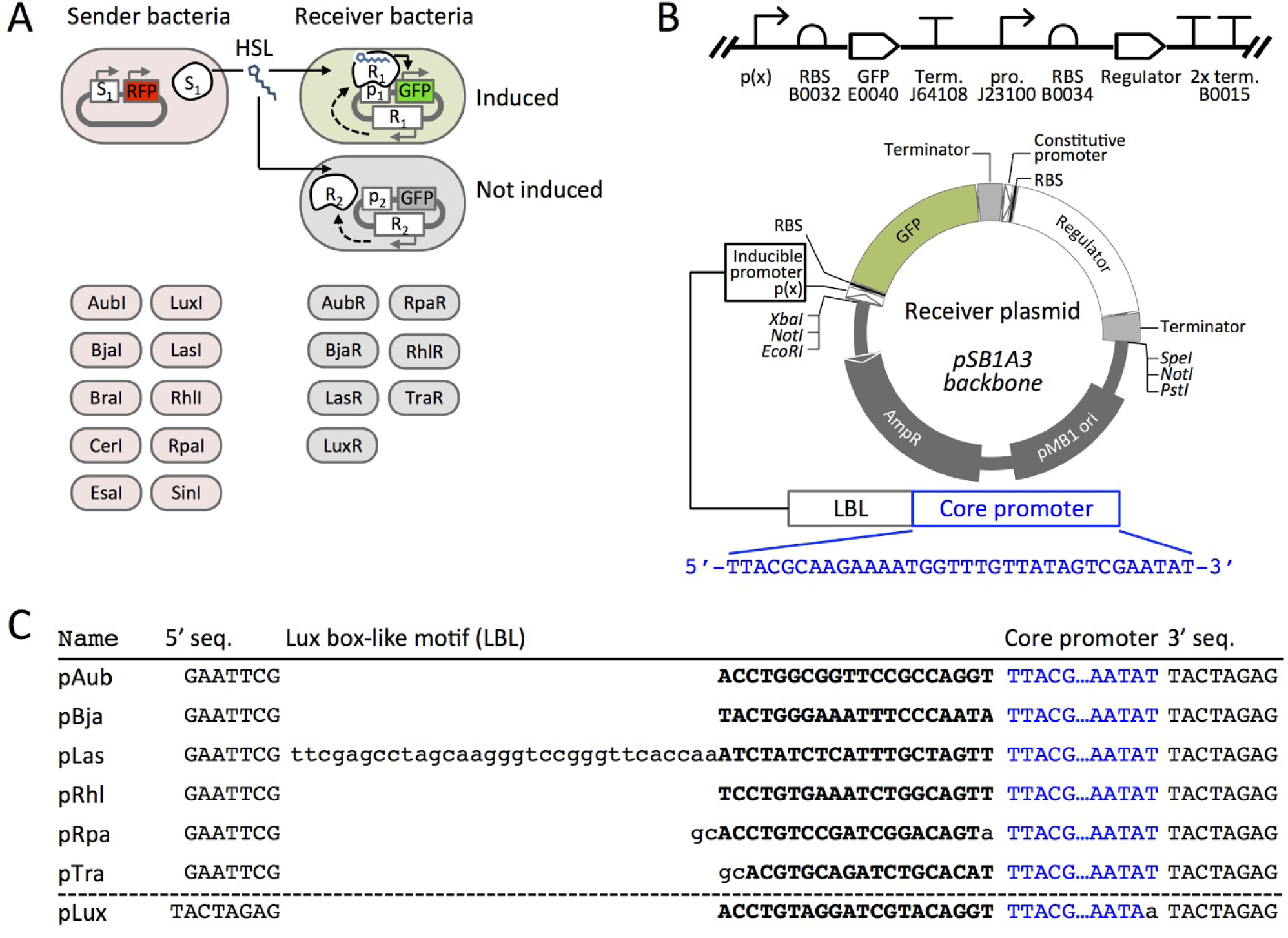
Scheme to determine HSL signaling orthogonality. **(A)** Sender plasmids encode an HSL synthase (S) and red fluorescent protein RFP, as previously described (Daer et al., 2018). HSL-inducible Receiver plasmids encode a constitutively-expressed regulator (R) and GFP controlled by the regulator’s cognate promoter (P). Each plasmid is transformed into a separate E. coli culture. **(B)** The SBOL diagram (top) and scaled circular map (bottom) represent the composition of six of the Receiver plasmid constructs: AubR, BjaR, LasR, RhlR, RpaR, and TraR. In the LuxR plasmid, the Regulator cassette is upstream of the inducible promoter and GFP. Codes indicate BioBrick ID numbers (e.g. B0032, etc.). RBS = ribosome binding site, Term. = transcriptional terminator, pro. = promoter. (C) Five inducible promoters (pAub - pTra) are synthetic hybrid sequences that each contain a Lux box-like motif followed by a Regulator-binding site (uppercase bold characters). pLux is from Canton et al. (Canton et al., 2008). 5’ seq. = 5’ sequence, 3’ seq. = 3’ sequence.

Here we define orthogonal pairs as vector transformed E. coli that transmit HSL signals between one Sender and one Receiver without affecting signalling between other pathways. A Sender shows potential orthogonal signaling behavior when it induces one or more Receivers but not others (Figure 1A). A potentially orthogonal Receiver is activated by one or a select few HSL signals. Identifying orthogonal quorum sensing pairs is challenged by at least two factors: synthases with broad HSL profiles and regulators that show non-specific responses to HSLs. As we have discussed in our previous report (Daer et al., 2018), many HSL synthases generate a combination of HSL molecules when the enzymes are heterologously expressed in E. coli. Thus, Senders that express these enzymes might promiscuously induce several different Receivers. We describe the behavior of a library of HSL-inducible regulators that respond to a diverse array of synthetic and synthase-produced HSLs, with some synthase-regulator pairs showing partial or complete orthogonality.

## Results and Discussion

### Design, Construction and Validation of Receiver Plasmids

We constructed a set of Receiver plasmids that encoded HSL-inducible green fluorescent protein (GFP) (Figure 1B). Each constitutively expresses a regulator protein, a transcription factor that gains affinity for specific promoter sequences when the protein’s binding pocket is occupied by an HSL ligand. The LuxR cassette used in our previous work (Daer et al., 2018) included a constitutively expressed Lux regulator gene upstream of an inducible GFP reporter. To avoid potential readthrough from the regulator gene into the GFP reporter in other constructs, we designed a new cassette where the inducible reporter was placed upstream (5’) of the constitutively expressed regulator. Also in the new design, we use a hybrid promoter topology (Figure 1C) inspired by the work of Scott et al., where HSL-regulated promoters include a lux-box like DNA binding domain upstream of a minimal promoter (Scott and Hasty, 2016). Each Lux box-like motif (LBL) acts as a target site for a corresponding HSL-bound regulator that helps initiate transcription from the weak downstream promoter.

We validated the function of each Receiver plasmid by measuring GFP expression over time from E. coli transformants that had been exposed to different concentrations of synthetic HSL compounds. To enable comparison across datasets, we included a constitutive GFP-expressing plasmid Ctrl-GFP. This plasmid (ptrc99a) included a trc promoter followed by a GFP ORF and an rrnB terminator. Overall, we observed dose-dependent responses for five of the Receiver plasmids: AubR, BjaR, LasR, LuxR, and TraR (Figure 2).

**Figure 2.**
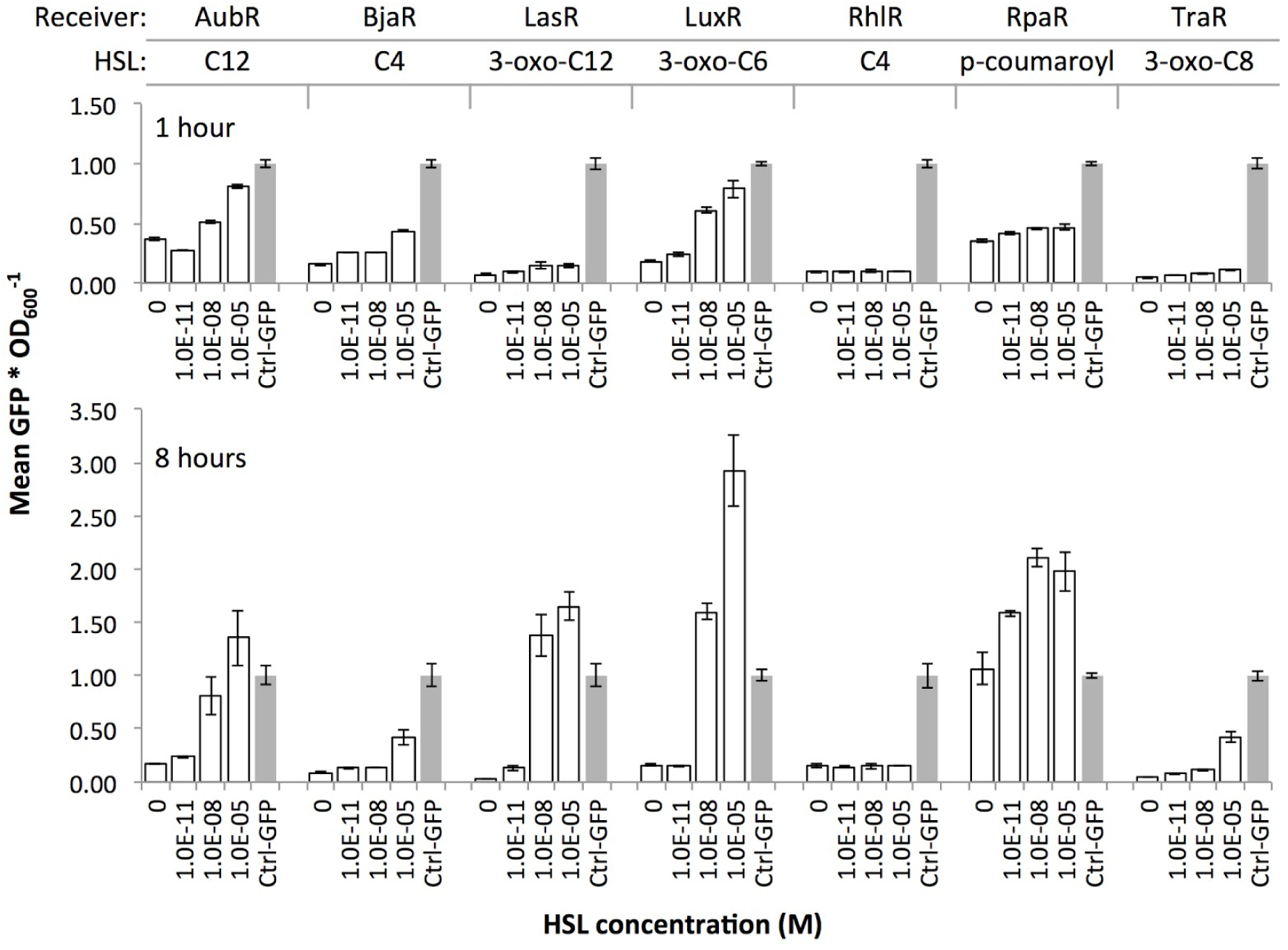
Induction of Receiver constructs with cognate HSL ligands. Receiver cells were grown in a 96-well plate in liquid media supplemented with zero, 1.0E-11, 1.0E-08, or 1.0E-05 of HSL compound. Constitutive GFP-expressing cells (Ctrl-GFP) were used as a positive control in each assay. Individual values (GFP signal / OD_600_) within each set were normalized by the mean Ctrl-GFP value. Bars = mean GFP signal / OD_600_ of three wells, error bars = standard deviation.

The data confirmed HSL-induced responses for all but two Receivers: RhlR and RpaR (Figure 2). The RhlR regulator failed to induce GFP at any concentration of its cognate HSL (C4). We observed the same outcome in a solid agar plate assay where RhlI Sender cells were plated near RhlR Receiver cells (data not shown). The RpaR Receiver showed dose-dependent induction at the 1 hour time point. However, at 8 hours all samples including the zero HSL negative control showed GFP signals at or above Ctrl-GFP, which suggested very leaky expression from the pRpa hybrid promoter. We omitted this regulator because the time frame for HSL dose-dependent induction might be too short to collect interpretable data with our assays. We included five of the Receivers, AubR, BjaR, LuxR, LasR, and TraR, in subsequent experiments.

### Inductions With Synthetic HSLs Revealed Distinct HSL-Response Profiles

To explore the specificity of the validated Receivers to HSLs, we measured the responses of the Receivers to various synthetic HSLs. Many natural regulators are known to respond to a single HSL within their native bacterial strain (Davis et al., 2015). These narrow-range signalling systems include Bja which generates a short chain HSL (4 carbons), and Tra, Aub, and Las which generate long chain HSLs (8, 12, and 12 carbons, respectively). Other systems are known to produce a broader range of two or more HSLs (Davis et al., 2015) such as Lux, which generates mid chain length HSLs (6 and 8 carbon). We hypothesized that if quorum sensing regulators evolved to respond only to the HSLs that are produced by each system, and that the recombinant Receiver regulators behave the same way, then AubR, BjaR, LasR, and TraR should respond to a single HSL, and LuxR should respond to multiple HSLs.

We grew each Receiver strain in liquid media that was treated with different concentrations of eight different HSL compounds with varying chain lengths (Figure 3). AubR was most sensitive to C12 HSL, as expected. This Receiver appeared to be much less responsive to 3-oxo C12, suggesting that the absence of the oxo group might contribute to HSL selectivity. AubR responds to a lesser extent to some similar chain length HSLs, such as C14 and C8, suggesting that there is some flexibility in the acyl tail binding pocket to accommodate other chain lengths at higher concentrations. However, AubR did not respond to short chain or large R group HSLs, indicating that the nonpolar acyl chain is an important factor in binding specificity.

**Figure 3.**
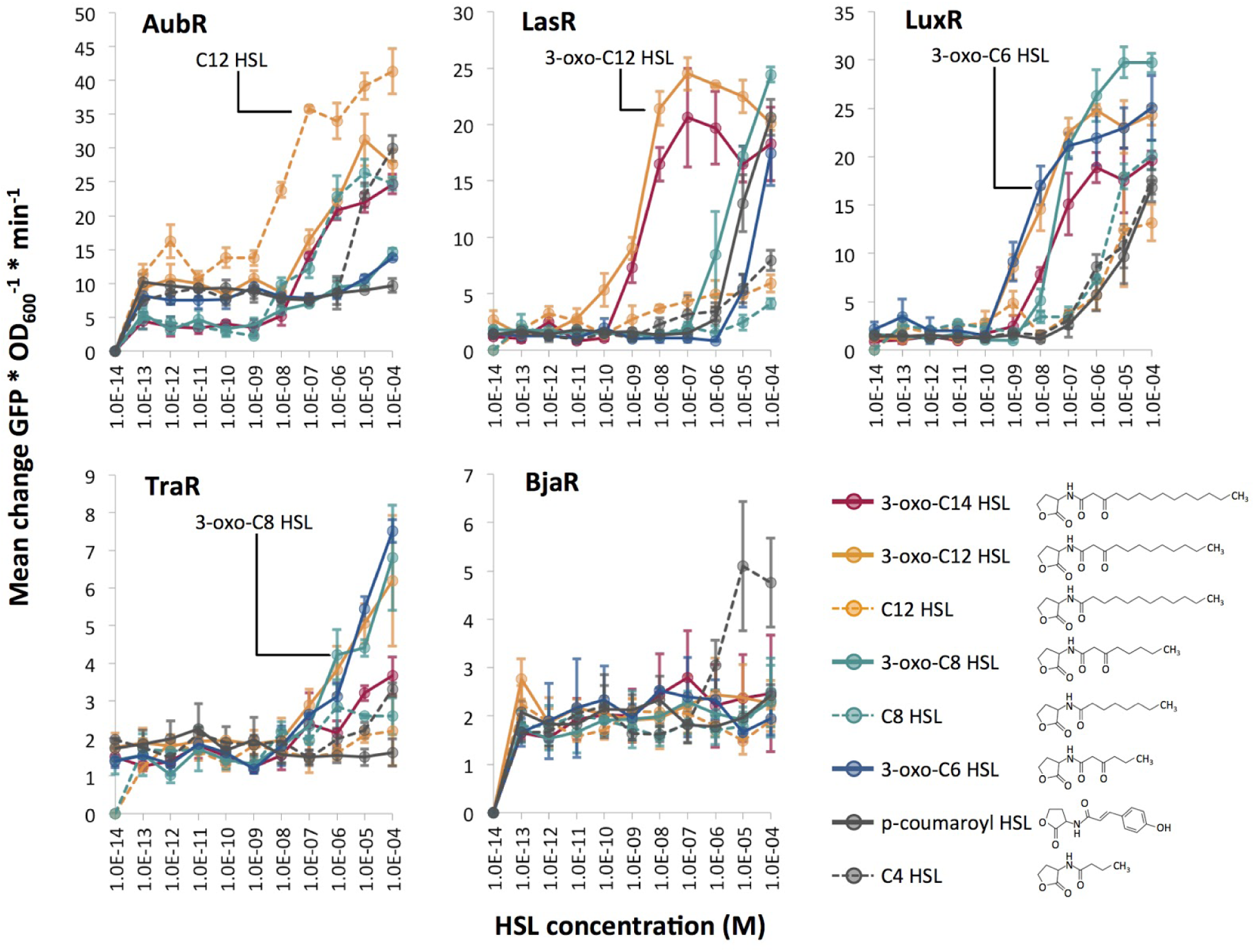
HSL dose response experiments for five Receivers. Synthetic HSLs (purchased from Sigma) were dissolved in DMSO. Receiver bacteria were cultured in the presence of 1.0E-14 to 1.0E-4 M HSL (x-axis) and measured for GFP expression over time. Each point represents the average (mean of three technical replicates) maximum rate of change of GFP/OD_600_ per minute. The cognate ligand for each regulator protein is labeled except for BjaR; isovaleryl HSL was not included in this assay.

LasR was similarly sensitive to 3-oxo-C12 and 3-oxo-C14 HSL. LasR might prefer the 3-oxo HSL side group more than HSLs of the same length without it, as seen when comparing induction with 3-oxo-C12 HSL to C12-HSL. LasR prefers acyl tail lengths close to its cognate ligand (12 carbon), but at higher concentrations, is able to accomodate shorter length HSLs. This binding flexibility has been previously demonstrated with biophysical data that showed LasR binding to a structurally diverse set of inducer mimics (Zou and Nair, 2009). This observation is consistent with our results which show that a variety of HSL chain lengths can activate LasR-mediated GFP expression (Figure 3).

LuxR showed the strongest response to 3-oxo-C6 and an intermediate response to 3-oxo-C8, which is consistent with its natural activity. These results agree with those reported by Canton et al. (Canton et al., 2008) which used the same synthetic LuxR system. However, our experiments included 3-oxo-C12 and 3-oxo-C14 HSLs, which induced LuxR to a similar degree as 3-oxo-C6 and 3-oxo-C8, respectively. There was little to no response to HSLs that lacked the oxo group, suggesting a strong preference for HSLs with this modification and less specificity in regard to length of the acyl chain.

TraR showed a similar preference for 3-oxo HSLs. The TraR Receiver was induced by all of the 3-oxo HSLs with mid-length acyl chains, C6, C8, and C12 (Figure 3) and not by the corresponding 3-oxo apo HSLs. Finally, BjaR was most sensitive to C4-HSL, as expected. BjaR did not respond to any mid or long chain HSLs, which agrees with previous work indicating that BjaR responds to the short-chain ligand isovaleryl HSL (Lindemann et al., 2011).

These results partially supported our hypothesis that AubR, BjaR, LasR, and TraR would be induced by a single HSL, and LuxR would respond to multiple HSLs. All of the Receivers showed strong responses to their cognate HSLs. BjaR behaved as a narrow range regulator and LuxR behaved as a broad range regulator as predicted. Contrary to our hypothesis, AubR, LasR, and TraR responded to a broad range of HSLs: five, five, and three HSLs respectively. Furthermore, LuxR appears to be as sensitive to longer chain length HSLs (C12, C14) as it is to C6 and C8.

Finally, LasR, TraR, and LuxR appear to prefer 3-oxo HSLs, whereas AubR prefers a ligand without the oxo group.

### Sender-Receiver Pairs Show Crosstalk and Orthogonality In Liquid and On Solid Cultures

Next we measured the responses of the Receivers to HSLs produced by Sender cells that expressed one of ten different synthases. The HSLs that are expected to be produced by the synthase orthologues cover a broader range of structural diversity (described in (Daer et al., 2018)) than the set of synthetic HSLs used in our previous induction assay (Figure 3). E. coli were transformed with a Receiver plasmid or a an HSL-synthase expressing Sender plasmid and grown overnight in liquid media. The next day, the Sender cells were pelleted and the HSL-enriched supernatant was filtered to eliminate Sender cells. Receiver cells were diluted in fresh media and inoculated with 10% - 50% volumes of HSL-enriched supernatant (illustrated in Figure 4) and measured for GFP signal over time (Supplemental Figure S1).

**Figure 4.**
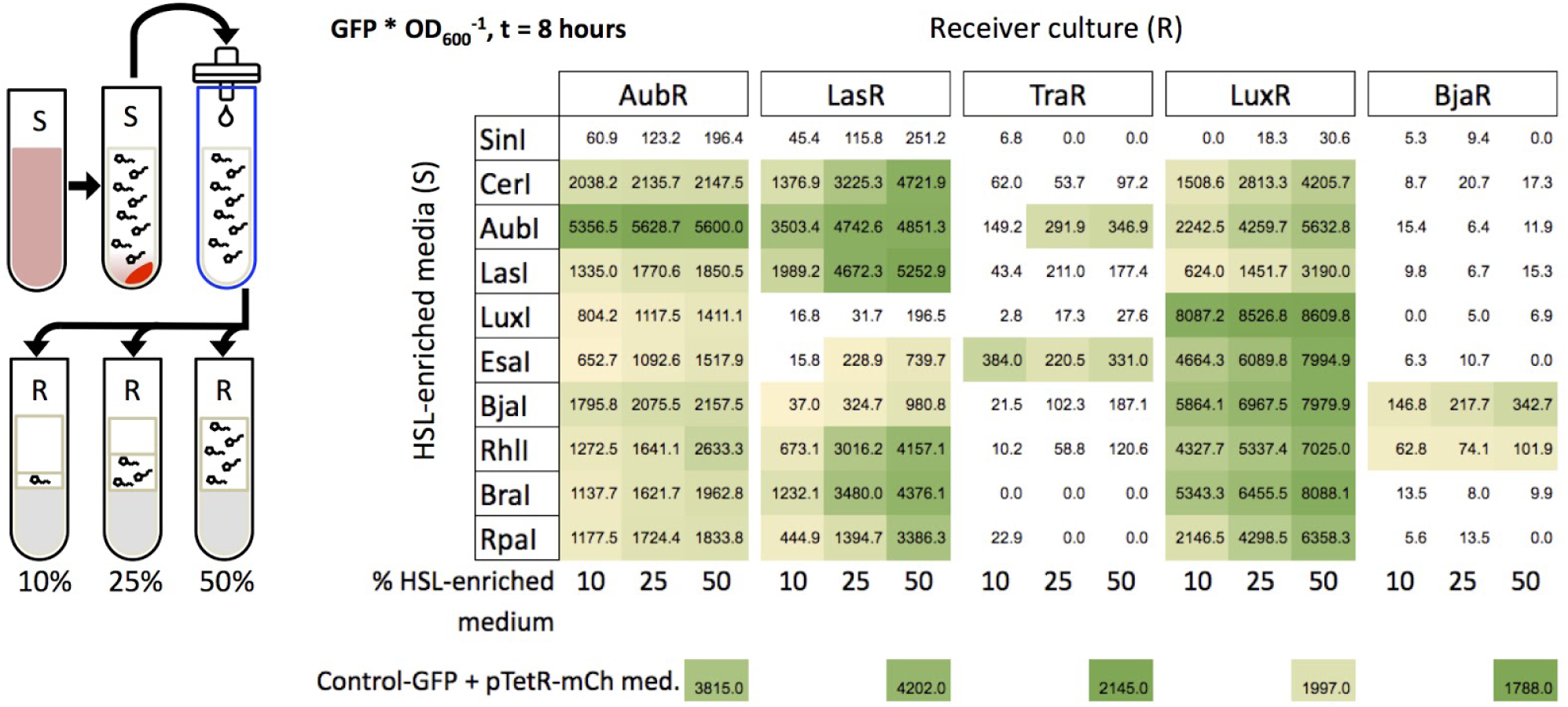
Induction of Receivers with HSL-enriched media. Sender cells (S) were grown in liquid cultures to produce HSL-enriched LB. Media collection and filtration were performed as described in Daer et al. (Daer et al., 2018) HSL-enriched media were brought to a final concentration of 10%, 25%, or 50% in Receiver cultures (OD_600_ = 0.4). Each value in the grid represents the mean of the maximum GFP/OD_600_ from three wells after subtraction of background (negative control) at 8 hours of induction. The negative controls are each Receiver treated with mock-enriched media (50% volume) from cells expressing mCherry from an “empty” Sender plasmid (pTetR-mCh med.). The positive control constitutively expresses GFP and was treated with the same mock-enriched media (Control-GFP + pTetR-mCh med.). Yellow to green shading indicates samples that showed higher GFP/OD_600_ at t = 8 hours versus t = 0, and values that are significantly greater than the negative control (p < 0.05, Student’s one-tailed t-test).

To aid in interpreting the data, we sorted the synthases based on the highest carbon chain length(s) of HSL(s) they are known to generate, i.e. SinI (C18) to RpaI (C4) (Figure 4). Four of the synthases were derived from the same systems as the regulators in our library. These cognate synthases include AubI, LasI, LuxI, and BjaI. A Tra Sender (Moré et al., 1996) is not included in our library. Eight of the synthases are known to generate only a single HSL (RpaI, RhlI, BjaI, AubI, CerI, LasI, BraI, EsaI), while two are broad range synthases that generate two or more different HSLs (SinI, LuxI) (Davis et al., 2015). Since many natural synthases are predicted to produce a single HSL, we expected the outcome of this experiment with cellular HSLs to agree with the results from the previous induction experiments with synthetic HSLs. However, this prediction assumes that sufficient levels of HSL metabolism substrates, specific acyl carrier protein species or coenzyme A, are available in E. coli BL21.

Overall, HSL-enriched media from each Synthase induced the strongest expression of GFP from its cognate Receiver compared to the other nine Synthases in each experiment. The strongest agreements with our prediction were observed for LasI and LuxI. LasI was the strongest inducer of LasR (Figure 4), indicating that LasI produces 3-oxo-C12-HSL as expected. Furthermore, AubR and LuxR which responded to synthetic 3-oxo-C12-HSL (Figure 3), were also induced by LasI. LuxI, which is known to produce 3-oxo-C6-HSL and C8-HSL, showed strong induction of LuxR and moderate induction of AubR. LuxR responded to both HSLs when tested individually (Figure 3). Since high concentrations of 3-oxo-C6-HSL did not induce AubR, LuxI may have stimulated AubR via C8-HSL.

Other cognate synthases showed evidence of non-specific HSL production. AubI, which we expected to generate C12 HSL, was the strongest inducer of AubR, strongly induced LasR and LuxR, and slightly induced TraR. This result suggests that in E. coli BL21 AubI produces more than one HSL in the range of 6 - 14 carbons, possibly with 3-oxo groups. BjaI which is known to generate isovaleryl HSL (Lindemann et al., 2011), was the strongest inducer of BjaR. We demonstrated that BjaR responded only to synthetic C4 HSL (Figure 3). Therefore, BjaI might produce isovaleryl HSL, C4 HSL or both. BjaI also induced the AubR, LasR, and LuxR Receivers (Figure 4), which responded weakly to synthetic C4 HSL (Figure 3). Low levels of LasR Receiver activation by BjaI suggest this synthase also produces small quantities of other HSLs, potentially in the C6 - C12 range.

One of the non-cognate synthases induced Receivers in a manner that was consistent with its HSL profile. EsaI, which is known to synthesize 3-oxo-C6 HSL (Beck von Bodman and Farrand, 1995), showed strong induction of TraR and LuxR, and weaker induction of AubR and LasR compared to the other synthases. Experiments with synthetic HSLs showed a similar trend, where TraR and LuxR were the very sensitive to 3-oxo-C6-HSL, and AubR and LasR had a relatively weaker response (Figure 3). These results suggest that EsaI produces the expected HSL. SinI, which produces C8, C12, 3-oxo-C14, 3-oxo-C16, and C18 HSLs (Marketon et al., 2002) showed no significant levels of Receiver activation. This enzyme may be intrinsically inefficient, or produce mostly longer acyl-chain HSLs that do not stimulate the five Receivers in our panel.

The other non-cognate synthases appeared to stimulate off-target Receivers. CerI is known to produce 7,8-cis-N-(tetradecenoyl)-HSL, a monounsaturated C14 HSL with a cis double bond between carbons 7 and 8. Strong induction of AubR by CerI-supernatant (Figure 4) suggests that promiscuity of the binding site for acyl chains of slightly different lengths might allow AubR to respond to C14 HSL. Alternatively, CerI might produce a broader range of HSLs than expected since CerI activated LasR and LuxR, which were sensitive to 3-oxo HSLs (Figure 3). RhlI, a C4 HSL generating synthase, induced every receiver except TraR (Figure 4). AubR, BjaR, and LuxR might be stimulated by C4 from RhlI since these Receivers were activated by high levels of synthetic C4-HSL. However, LasR and TraR were insensitive to synthetic C4, thus RhlI might produce additional HSLs that activate these Receivers. RpaI encodes a synthase that produces p-coumaroyl HSL, an aryl-HSL with a phenol R group (Schaefer et al., 2008). LasR and LuxR were activated by supernatant media from RpaI (Figure 4), which is consistent with the moderate stimulation of these Receivers by synthetic p-coumaroyl HSL (Figure 3). However, RpaI-enriched media also induced AubR (Figure 4), which suggests that RpaI produces one of the larger HSLs (e.g. C12 HSL). BraI, another low molecular weight aryl-HSL synthase, had a Receiver induction profile that was very similar to that of RpaI. BraI is known to produce cinnamoyl-HSL with a phenyl R group (Ahlgren et al., 2011). Activation of AubR in our assays suggests that BraI also produces at least one larger acyl-HSL.

Although several of the Senders and Receivers showed varying degrees of cross-signaling, no single Sender showed significant induction of all five Receivers, and just two Receivers, AubR and LuxR, showed sensitivity to nine Senders in all HSL-media dilutions. Therefore, orthogonal behavior might be achieved by using carefully-selected pairs of Senders and Receivers, as we describe later in this report. Having found evidence of orthogonality using fixed amounts of HSLs, we were interested in determining if the same results could be observed under conditions where Receivers were exposed to continuous HSL production over time by live Sender cells.

Sender and Receiver cells were plated on solid LB agar as shown in Figure 5. Sender cells were streaked near a rectangular lawn of Receiver cells, as previously described (Daer et al., 2018). This configuration allowed Receivers to be exposed to HSLs as the molecules diffused through the medium. Thus, we could monitor relative induction distance over time by using 3D renderings generated from 2D images of GFP signal over the surface of each plate (Figure 5). The results from the agar based assays mostly agreed with those from the liquid culture experiments with a few key exceptions. While SinI showed no significant induction of any Receiver in the liquid culture assays, the results from agar cultures showed modest induction of AubR and LasR. SinI is known to produce C8, C12, 3-oxo-C14, 3-oxo-C16, and C18 HSLs (Marketon et al., 2002). C12 HSL may induce AubR and 3-oxo-C14 might drive activation of LasR. Also contrary to the results from liquid culture experiments, TraR was not strongly induced by AubI, and LasR was not strongly induced by EsaI, RhlI, and BraI (Figure 5). In general, this difference is a loss of weak or off-target induction that was observed in the previous liquid culture assays.

**Figure 5.**
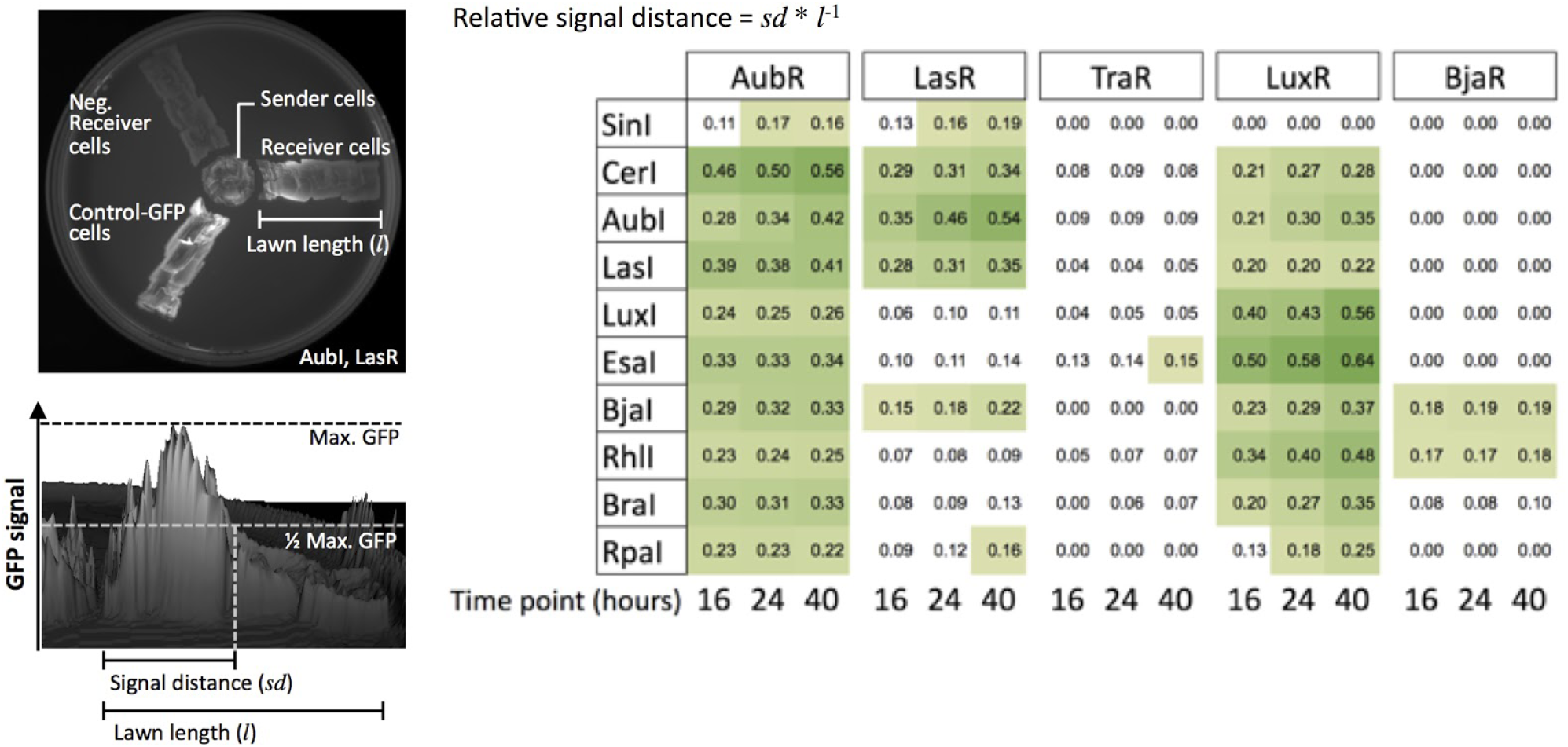
Induction of Receiver cells by Sender cells grown on solid agar. The configuration of the agar plate assay for LasI (Sender) and LasR (Receiver) is shown (upper left). Genepix software was used to render the 3D image and quantify GFP signal from the Receiver cells (lower left). The color-scaled grid shows values (one trial) of relative GFP signal distance over a rectangular region of plated Receiver cells for every Sender/ Receiver pair tested. Yellow to green squares are values of 0.15 or higher.

Two key differences in experimental conditions might underlie these outcomes. Gains in Receiver sensitivity (i.e. SinI activation of AubR and LasR) may have been supported by constant production of HSLs by live Senders on agar, whereas only fixed quantities of HSLs were present in the liquid culture experiments. Additionally, in the liquid culture experiments GFP/OD_600_ decreased over time in some samples, suggesting that dilution of GFP per cell by cell division or degradation might dampen GFP signal output. Losses of TraR and LasR Receiver induction by AubI, EsaI, RhlI, and BraI may have been driven by constrained diffusion of HSLs in agar, favoring the induction of Receivers by the predominant HSLs rather than secondary products of the synthases.

### Identification of Orthogonal Pathway Sets

We used the data from the liquid and agar culture experiments to identify orthogonal pathway sets that might be useful for the development of layered logic gates. Here, we define an orthogonal pathway set as two (or more) distinct Receivers that all become activated only in the presence of two (or more) distinct Senders. This can be accomplished when HSL signalling from each Sender activates only one Receiver within the set. Both the liquid culture assays and agar assays indicated a broad range of crosstalk, albeit to a lesser extent for the agar assay data. Still, we were able to identify two potentially orthogonal quorum sensing pathway sets that were supported by data from both experiments.

The results from the liquid culture induction experiments are summarized in Supplemental Figure S2 as a Sender-Receiver induction chart. Although the chart indicates a wide range of crosstalk, we could still identify two sets of Sender/Receiver pairs that might function orthogonally under specific conditions. BjaI/BjaR and EsaI/TraR (set 1) should function in parallel without crosstalk at high and low concentrations of HSLs. EsaI/TraR and LasI/LasR (set 2) signalling could be carried out in parallel, as long as HSL concentrations are kept at low levels (10% HSL-enriched media), but are sufficient to induce the on-target Receivers.

From our agar plate induction results, we can predict several additional orthogonal sets of Sender/Receiver pairs (Supplemental Figure S3). RhlI/BjaR and LasI/LasR (set 3), and SinI/LasR and RhlI/LuxR (set 4) exhibit weak crosstalk for one of the two Receivers. This crosstalk is well below the induction strength of the on-target signaling. Our data also suggest that three Sender/Receiver pairs, LasI/LasR and BjaI/BjaR and EsaI/TraR (set 5), can be used in parallel.

Crosstalk could be mitigated by increasing the distance between Senders and Receivers to disfavor weak induction of off-target Receivers. This idea is supported by the streak plate experiments where weak inducers showed GFP over shorter distances. Figure 6 shows results from a qualitative spot agar assay to verify Sender-Receiver signalling in the predicted orthogonal sets. The high level of off-target induction observed for LasI/LuxR (set 4) underscores the importance of tuning the sensitivity of signaling in order to accomplish complete orthogonality. All together, we can predict two strong candidate sets of engineered quorum sensing pathways that function without crosstalk in liquid and solid agar media: BjaI/BjaR and EsaI/TraR and EsaI/TraR and LasI/LasR.

**Figure 6.**
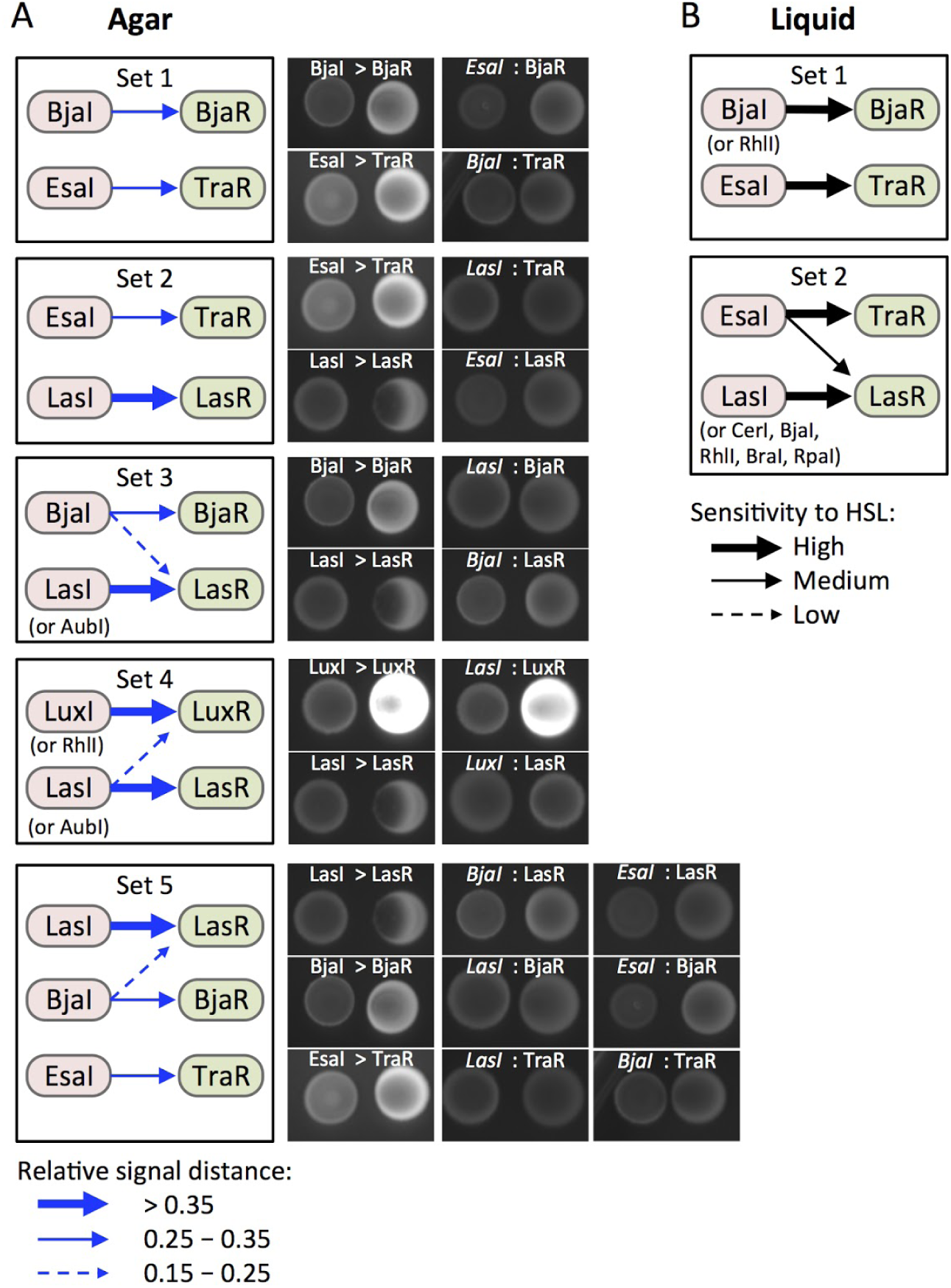
Identification and validation of candidate orthogonal pairs. (**A**) Boxes (Sets 1 - 6) show Sender/Receiver pairs for which HSL signalling is predicted to show no or minimal crosstalk based data from the agar culture assays (Figure 5). Induction strength (blue arrows) is represented by relative induction distance values. Strong, medium, and weak induction are defined by the value thresholds shown in the legend (bottom left). Labeled images show fluorescent protein signal from 5 μL spots of Sender (left, RFP) and Receiver (right, GFP) cells after 24 hours of growth on LB agar. (**B**) Boxes (Sets 1 and 2) show signaling pairs for which results from liquid culture assays (Figure 4) also suggest orthogonality. High, medium, and low sensitivity (black arrows) represent inductions where 10%, 25%, or 50% HSL-enriched medium increased GFP signal from a Receiver compared to mock-enriched medium.

## Conclusions

We used a decoupled HSL synthase and regulator-reporter system to investigate signalling between quorum sensing pathway modules. We present evidence that several regulators show a broader range of sensitivity than expected to different HSLs in the context of an engineered operon (Receiver plasmids) and in a non-native host cell, E. coli BL21. Our results also provide evidence that certain synthases produce more bioactive HSL species than expected when they are overexpressed in E. coli. Our characterization of Bja and Aub components in a non-native host contributes new information to the growing body of quorum sensing research. Finally, we used a systematic investigation of quorum sensing crosstalk to predict orthogonal sets of Sender/Receiver pairs for bioengineering applications.

Our results suggest that crosstalk can arise from regulator promiscuity through interactions with multiple HSL ligands, and non-specific generation of HSLs by recombinantly expressed synthases. Our current data for LuxR recapitulates results from previous work where we demonstrated that a LuxR Receiver responded to a wide range of HSLs from Sender strains (Daer et al., 2018). Four new Receivers, AubR, LasR, TraR, BjaR, were expected to respond specifically to the HSL that is associated with each ortholog in nature. However, AubR and LasR responded to multiple synthetic and cellular HSLs. LasR’s capacity to bind to a diverse set of HSLs and other small molecules is supported by previous structural and biochemical studies (Borlee et al., 2010; Zou and Nair, 2009). We are the first to report a broad HSL induction profile for the Aub regulator. Our work also suggests that certain synthases produce a broader range of HSLs than expected. Eight of the ten synthases were expected to generate a single HSL species. AubI, BjaI, CerI, RhlI, RpaI, and BraI induced Receivers in a manner that was inconsistent with the synthases’ known HSL profiles. Our results for RpaI are consistent with a previous study that showed substrate-dependent production of a variety of HSLs by RpaI (Kang et al., 2015). The data in our current report is the first evidence of the generation of multiple bioactive HSLs from AubI, BjaI, CerI, RhlI, and BraI.

It is important to consider the influence of the host’s intracellular environment on the behavior of synthases and regulators. Our work investigated regulators in a non-native host, E. coli BL21, which only has one known Lux-like regulator protein, SdiA (Lindsay and Ahmer, 2005), and no known HSL synthase enzymes. Regulators that showed little or no expression GFP in the presence of HSLs (i.e. TraR, BjaR, and RhlR) might be incompatible with E. coli specific transcription factors, or their associated lux-box like sequences may not function well with the weak core promoter. Furthermore, E. coli SdiA which binds 3-oxo-C6 and 3-oxo-C8-HSLs (Lindsay and Ahmer, 2005) might sequester HSLs and dampen regulator activity. In our experiments, the RpaR Receiver showed HSL-independent GFP expression, which suggests that factors in E. coli contributed to hyperactivity. SdiA is known to bind the Rhl promoter (Lindsay and Ahmer, 2005) so it is possible that SdiA can interact with pRpa. This idea would need to be tested directly with a pRpa-GFP reporter that lacks RpaR. Another plausible explanation for HSL-independent GFP expression from the RpaR Receiver is hyperactivity of the regulator. High concentrations of overexpressed RpaR might favor dimerization, promoter binding, and GFP transcription. Incorporation of a conserved tryptophan via a gain-of-function mutation is required for TraR activity in E. coli (Scott and Hasty, 2016), suggesting that less robust regulators lack intrinsic transcription factor recruitment activity. This suggests a strategy of directed mutagenesis of weak or non-functional regulators may enhance their functionality in heterologous hosts.

It is important to note that in our engineered Sender/Receiver system, no more than one type of regulator was co-expressed in a single cell. Certain regulator proteins can bind to off-target lux box promoter motifs when two or more are co-expressed in a single cell (Scott and Hasty, 2016). Therefore, the orthogonality we describe here might require each Receiver to be expressed by a single transformed strain. This type of platform is relevant and useful for coordinating cellular phenotypes and dynamic behaviors in synthetic microbial consortia. For instance, intercellular quorum sensing control has been demonstrated in liquid media where co-cultured strains carried out self-regulation (Scott et al., 2017) or cross-regulation (Balagaddé et al., 2008) of cell growth. Multistrain regulation has also been demonstrated on solid media where HSL-signalling colonies were used to generate XOR logic gates (Tamsir et al., 2010). Recently, quorum sensing systems were used to execute information transfer between bacteria within a mammalian gut (Kim et al., 2018). Teams of undergraduate scientists in the International Genetically Engineered Machines Competition (iGEM) have demonstrated creative use of multi-strain quorum sensing systems for a variety of applications such as biosensors, circuits, and computational modeling. Team Braunschweig 2013 designed a three-regulator system to cross-regulate the expression of antibiotic resistance genes to stabilize population diversity (Team:Braunschweig - 2013.igem.org). Team ETH Zurich 2013 used Sender and Receiver strains plated on agar to mimic a popular electronic puzzle game called Minesweeper (Team:ETH Zurich - 2013.igem.org). In 2018, Team ECUST used a quorum sensing regulator in E. coli to detect HSLs from A. ferrooxidans, a rust-producing bacterium (Team:ECUST - 2018.igem.org). The new quorum sensing tools reported in our work could advance further development of engineered microbial consortia.

## Materials and Methods

### Construction of Plasmid DNA

All sequences are available at https://benchling.com/hayneslab/f/UWzcp3nk-quorum-sensing-collection/. Gene cassettes were designed in CLC Main (Qiagen) and synthesized as g-blocks by IDT. G-blocks were PCR amplified using Forward primer 5’-TTATTCGAATTCGCGGCCGCTTCTAGAG and Reverse primer 5’-GGATTTCTGCAGCGGCCGCTACTAGTA with the following conditions: in a 50 μL reaction: 10 μL 5x HF buffer (New England Biolabs, NEB #E0553L), 2 μL 10 mM dNTP mix (NEB #N0447L), 1 μL 100 μM forward and reverse primer, 25 ng g block template, and 0.5 μL phusion polymerase. Reactions were cycles as follows: 45 seconds at 98°C, 25 cycles of: 10 seconds at 98°C, 20 seconds at 68°C, 1:15 at 72°C, followed by a final extension for 5:00 at 72°C. PCR products were column purified (Qiagen PCR cleanup kit #28104), and 2 μg of DNA was digested with EcoRI-HF and PstI-HF (NEB) with the manufacturer’s recommended conditions, followed by another PCR cleanup (Qiagen #28104). 6 μg of backbone vector pSB1A3 was digested with EcoRI-HF and PstI-HF (NEB), electrophoresed on a 1% agarose gel for 45 minutes, and gel extracted (Sigma #NA1111-1KT). Purified, digested insert and vector were ligated with the following conditions 1 μL 10X T4 DNA ligase buffer (NEB), 50 ng vector, 150 ng insert, 0.5 μL T4 DNA ligase (NEB) in 10 μL total for 10 minutes at room temperature. The ligation was transformed into chemically competent BL21 (NEB #C2530H) with the following conditions: 5 μL of ligation into 50 μL cells on ice for 5 minutes, followed by 45 seconds at 42°C, followed by 2 minutes on ice. 350 μL SOC was then added to the cells and incubated at 37°C, 220 RPM for 30 minutes. Cells were plated on LB/agar plate supplemented with 100 μg/mL ampicillin and grown overnight at 37°C. Purified plasmids were sanger sequenced for verification of insert before use.

### Agar Plate Inductions

All agar plate inductions were carried out on LB agar (Miller) supplemented with 100 μg/mL ampicillin. All cells were transformed into BL21 cells (New England Biolabs), plated on LB/agar plates, and grown overnight at 37°C to produce single colonies. Colonies from these plates were picked and streaked generously to fill in guided lines on fresh LB/agar plates using a pipette tip that was sterilely bent 90 degrees for streaking bacteria. Templates were printed and taped to the bottom of agar plates to help ensure consistency (0.75 cm radius circle with 3 cm long x 1 cm wide curved rectangles 0.2 cm from circle equally spaced around the circle). Plates were imaged on a Pxi4 imager at their respective time points using the 488 nm blot setting with automatic image focus and exposure. 2D images were converted to 3D using the Pxi4 software. Images were saved as TIFFs and induction distances were analyzed on Adobe Illustrator CC 2015.

### Cell Culture

Generation of cell synthesized HSLs and receiver cultures: 3.0 mL LB supplemented with 100 μg/mL ampicillin was inoculated with a single colony of BL21 (New England Biolabs) transformed with Sender, ptrc99a-(GFP), Receiver, or negative sender vector. Cells were grown in aerated 15 mL conical tubes overnight at 220 RPM at 37°C. All sender cell supernatants and negative sender supernatant controls were spun at 4200 RPM for 4:00 minutes before filtering supernatant through a 0.22 μm cellulose filter. All receiver cells were spun at 1000 RCF for 5 minutes to pellet cells, followed by removing the supernatant and then diluting the cells in fresh LB supplemented with 100 μg/mL ampicillin to OD_600_ =0.8.

### Inductions With HSL-enriched Media

96 well plate with lid (Sigma #CLS3603) contained 300 μL per well and scanned using a Biotek Synergy H1 Microplate Reader with the following settings: Incubate at 37°C with constant, maximum shaking (orbital frequency: 807 cpm). Every 10 minutes, read A600, and GFP fluorescence (485 nm excitation, 515 nm emission) for a total of 8 hours. Additional plate settings: Autoscale GFP, scale to low wells (Receiver + 50% negative sender wells (3 wells)), 200 scale value. All initial OD_600_ were 0.4 after dilution with sender supernatant. Negative sender sample consisted of 150 μL of negative sender cells resuspended in fresh LB/amp and 150 μL of filtered negative sender supernatant. GFP+ controls consisted of 150 μL GFP+ cells diluted in fresh LB/amp and 150 μL of filtered negative sender supernatant. Cells induced with 10% supernatant consisted of 150 μL of receiver cells diluted in fresh LB/amp, 30 μL filtered sender supernatant, 120 μL filtered negative sender supernatant. Cells induced with 25% supernatant consisted of 150 μL of receiver cells in fresh diluted LB/amp, 75 μL filtered sender supernatant, and 75 μL filtered negative sender supernatant. Cells induced with 50% supernatant consisted of 150 μL of receiver cells diluted in fresh LB/amp, and 150 μL filtered sender supernatant.

### Inductions With Synthetic HSLs

All HSLs were dissolved at a stock concentration in DMSO and serially diluted into DMSO. Each well consisted of 150 μL receiver cells diluted in fresh LB (OD_600_ = 0.8) supplemented with 100 ug/mL ampicillin, 3 μL HSL dissolved in DMSO, and 147 μL of negative sender supernatant (Final OD_600_ = 0.4).

### Spot Cultures on Solid Agar

Single colonies of senders and receivers were picked and grown overnight in 3 mL LB supplemented with 100 μg/mL ampicillin at 37°C and 220 RPM. Receivers were diluted 1:10 in fresh LB/amp, and 5 μL was dispensed by micropipette onto an LB/agar plate supplemented with 100 ug/mL ampicillin. 5 μL of undiluted Sender culture was applied 2.5 cm away from the center of the Receiver spot, leaving 0.5 cm between spots. The spots were allowed to air dry for 5 minutes, then incubated overnight at 37°C. Imaging of GFP was performed after 24 hours.

## Author Contributions

SJT: plasmid design and construct assembly, design and execution of liquid culture and streaked/ spotted agar plate inductions, GFP imaging; CLS: assistance with plasmid design and construct assembly; BL, CLS: synthetic HSL inductions; AM, CLS: sender supernatant inductions; CC, CLS, and XL: streaked agar plate inductions; SJT, KAH: project design, data analysis, manuscript writing; KAH: figures.

## Acknowledgements

This project was supported by a Kern Entrepreneurial Engineering Network (KEEN) student research grant (Arizona State University). B. Lopez, C. Smith, and A. Mani were supported by the Fulton Undergraduate Research Initiative (FURI). X. Livingstone was supported by the Western Alliance to Expand Student Opportunities (WAESO, NSF Grant HRD 1401190). S. Tekel was supported by the School of Biological and Health Systems Engineering (SBHSE). DNA synthesis and primers were generously provided by Integrated DNA Technologies (IDT) as support for the 2017 International Genetically Engineered Machines (iGEM) competition. We would like to acknowledge Cynthia Gallaway for assistance with managing the KEEN grant. We would also like to acknowledge Kylie Standage-Beier for constructive comments for editing the manuscript.

