## Supporting Information for "Engineered orthogonal quorum sensing systems for synthetic gene regulation"

S1 Table: iGEM Registry IDs for sequences and constructs used in this study

S1 Figure: Induction over time of receivers with sender supernatant

S2 Figure: Sender-Receiver induction map based on results from assays with cell-free, HSL-enriched liquid media

S3 Figure: Sender-Receiver induction map based on results from assays on solid agar

| **Part name** | **iGEM Registry part number** | **Contributing iGEM Team** |
| --- | --- | --- |
| LasR Receiver | BBa_K2357000 | iGEM17_Arizona _State |
| TraR Receiver | BBa_K2357028 | iGEM17_Arizona _State |
| AubR Receiver | BBa_K2357014 | iGEM17_Arizona _State |
| BjaR Receiver | BBa_K2357007 | iGEM17_Arizona _State |
| LuxR Receiver^(a)^ | BBa_K2357021 | iGEM17_Arizona _State |
| RpaI | BBa_K1421006 | iGEM14_CAU_China |
| BraI | BBa_K2033004 | iGEM16_Arizona_State |
| RhlI | BBa_C0170 | Antiquity |
| BjaI | BBa_K2033002 | iGEM16_Arizona_State |
| EsaI | BBa_K1670004 | iGEM15_Manchester-Graz |
| LuxI | BBa_C0161 | Antiquity |
| SinI | BBa_K2033008 | iGEM16_Arizona_State |
| AubI | BBa_K2033000 | iGEM16_Arizona_State |
| LasI | BBa_C0078 | Antiquity |
| CerI | BBa_K2033006 | iGEM16_Arizona_State |

**S1 Table.** iGEM Registry IDs for sequences and constructs used in this study. “Antiquity” indicates that no specific team is known to have contributed the DNA sequence to the Registry. Entries can be accessed at <http://parts.igem.org/>. (a) BBa_K2357021 is a derivative of BBa_F2620. An ORF encoding GFP-mut3b (BBa_E0040) was cloned downstream of BBa_F2620.

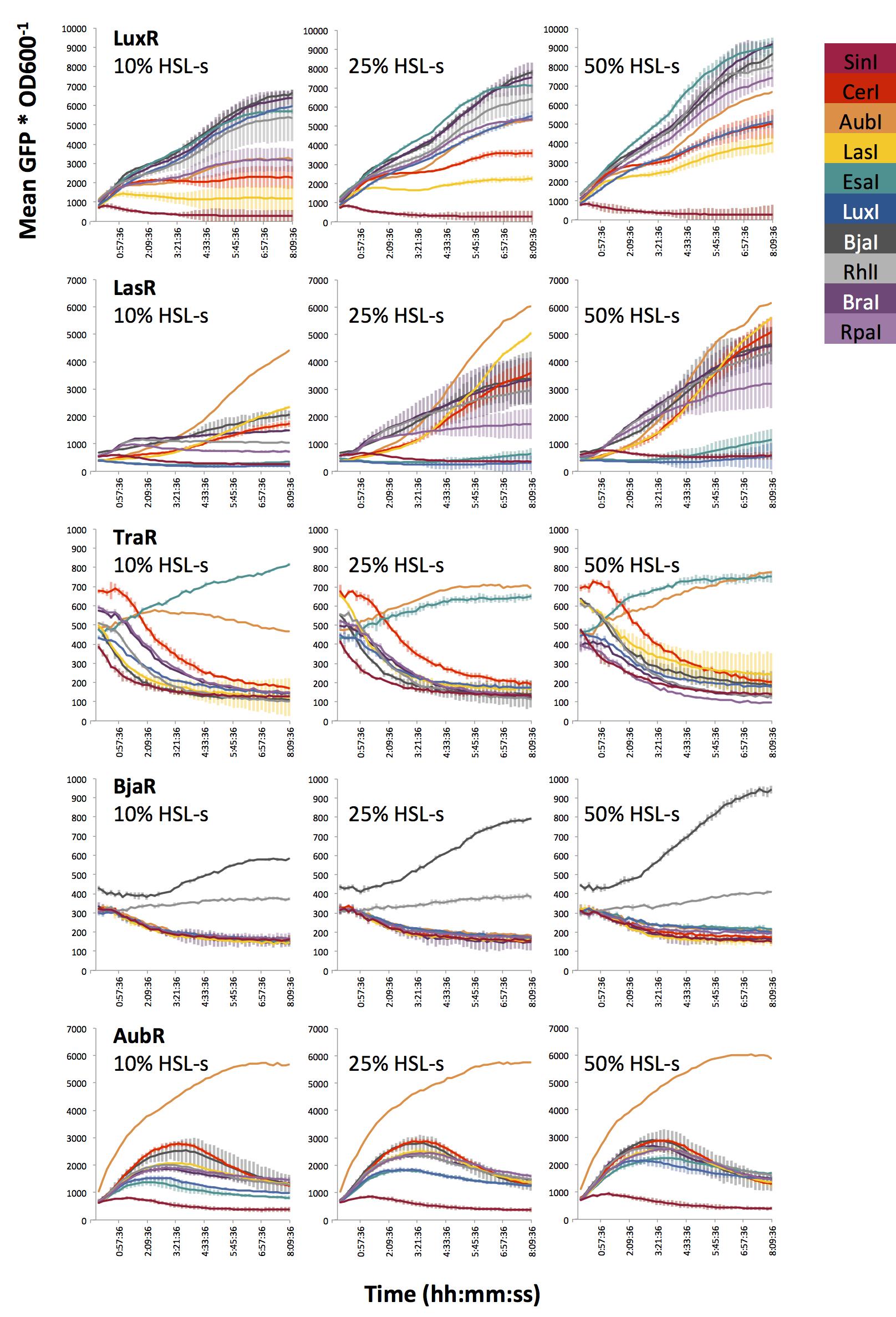

**Figure S1**. Induction over time of receivers with sender supernatant. Average GFP/OD_600_ over time. Receivers cells were diluted to OD_600_ = 0.4 in fresh LB and inoculated with filtered, HSL-enriched supernatant (HSL-s). Percent HSL is calculated based on the fraction of HSL-enriched supernatant in the total volume of treated Receiver culture (see Figure 4). Cells were incubated at 37℃ with constant orbital rotation with GFP (excitation 485, emission 515). GFP and OD_600_ were measured every 10 minutes over 8 hours. Error bars represent standard deviation of three wells.

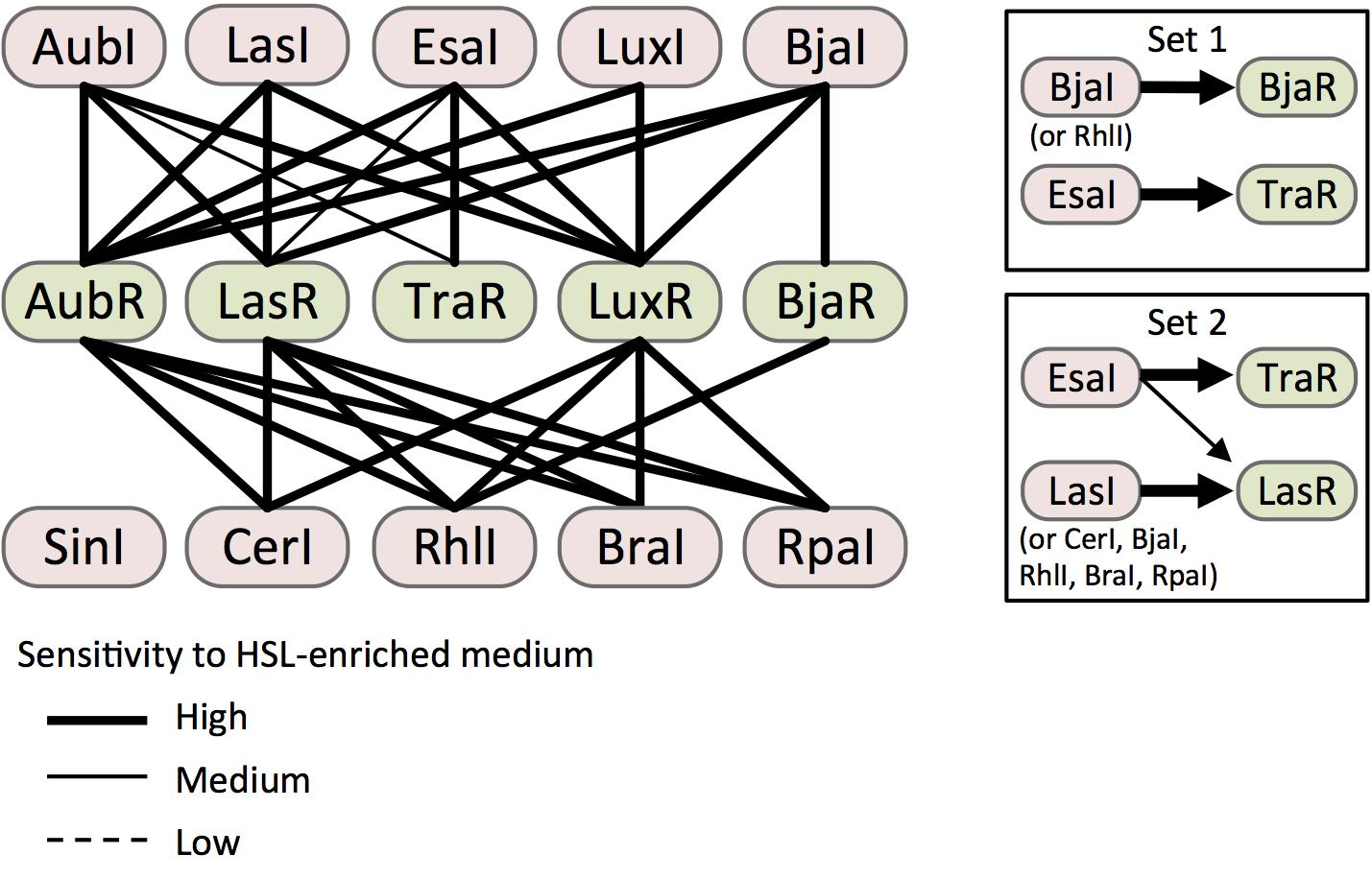

**Figure S2**. Sender-Receiver induction map based on results from assays with cell-free, HSL-enriched liquid media. Cells are represented as nodes (ovals) linked by edges that are formatted to represent the sensitivity of each Receiver (green) to the HSL-enriched supernatant produced by each Sender (pink). High, medium, and low sensitivity represent inductions where 10%, 25%, or 50% HSL-enriched medium significantly increased GFP signal from a Receiver compared to mock-enriched medium (*p* < 0.05, Student’s one-tailed t-test). Sets 1 and 2 (boxes) represent Sender/Receiver pairs that are expected to operate without crosstalk. Set 2 is predicted to exhibit no crosstalk at low HSL concentrations (i.e. 10% HSL-enriched medium).

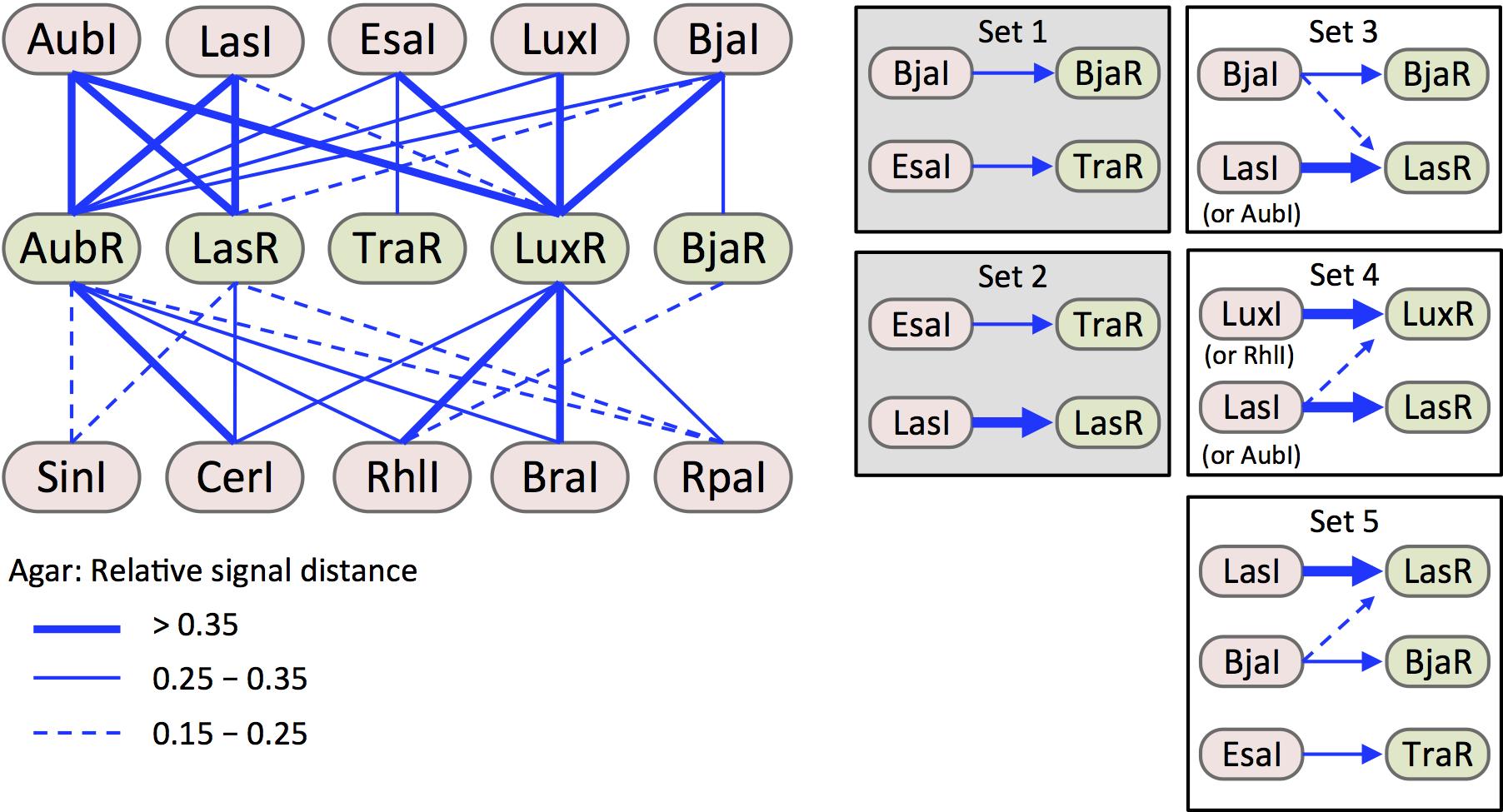

**Figure S3**. Sender-Receiver induction map based on results from assays on solid agar. As in Figure S2, Sender (pink) and Receiver (green) cells are represented as nodes (ovals) linked by edges that are formatted to represent the strength of induction. Here, induction strength is represented by relative induction distance across a lawn of Receiver cells, which was determined as described in Figure 5. Strong, medium, and low induction strengths, are classified by distance thresholds shown in the legend (bottom left). Shaded boxes indicate orthogonal sets that are also predicted by induction with HSL-enriched media in liquid cultures (sets 1 and 2 from Figure S2).
